# M1C IS NECESSARY FOR DARAXONRASIB RESISTANCE OF NSCLC KRAS(G12C) MUTANT CELLS

**DOI:** 10.64898/2026.06.20.733526

**Authors:** Shinkichi Takamori, Naoki Haratake, Kentaro Nonaka, Mai Moriya, Atrayee Bhattacharya, Tomoyoshi Takenaka, Tomoharu Yoshizumi, Mark D. Long, Donald Kufe

## Abstract

**Introduction:** The RAS(ON) multi-selective daraxonrasib (RMC-6236) inhibitor is effective in patients with NSCLC KRAS mutant cancers. Tolerance to daraxonrasib invariably develops by mechanisms that remain unclear. There is no known involvement of the M1C oncogenic protein in daraxonrasib resistance.

**Methods:** NSCLC H358 KRAS(G12C), H2122 KRAS(G12C) and patient derived MGH1112 KRAS(G12C) cells with acquired daraxonrasib resistance were investigated for M1C dependence in studies of SHP2, STAT1/3 and NF-KB activation, clonogenicity, and self-renewal capacity.

**Results:** We demonstrate that M1C is induced as a protective response in NSCLC KRAS(G12C) mutant cells treated with daraxonrasib. We report that M1C forms novel cell membrane-associated biomolecular condensates with the SHP2 protein tyrosine phosphatase in driving daraxonrasib resistance. M1C integrates SHP2 activation with induction of (i) oncostatin-m/gp130/STAT3 signaling, and (ii) the NF-κB-mediated epithelial-mesenchymal transition (EMT) pathway. The functional significance of this M1C-driven pathway is supported by the demonstration that targeting STAT3 and NF-κB reverses daraxonrasib resistance. Consistent with M1C dependence, we also show that targeting M1C is effective against daraxonrasib-resistant NSCLC KRAS mutant cell line and tumor models. In contrast, M1C drives sotorasib resistance by STAT1-mediated inflammatory signaling, demonstrating that M1C confers resistance to KRAS(G12C)-selective and RAS(ON) tri-complex inhibitors by noncongruent mechanisms.

**Conclusions:** These findings demonstrate that M1C is required for daraxonrasib tolerance and is a potential target for the treatment of patients with NSCLC KRAS(G12C) mutant tumors refractory to this agent.

## Introduction

Daraxonrasib is a RAS(ON) tri-complex inhibitor (TCI) that binds to active GTP-bound RAS and recruits cyclophilin A (CYPA) to block wild-type and mutant RAS(ON) signaling ^1, 2^. Daraxonrasib has substantial activity in patients with previously treated metastatic PDAC RAS mutant cancers ^3, 4^. Promising activity of daraxonrasib in patients with NSCLC RAS mutant tumors is being evaluated in the ongoing Phase 3 RASolve 301 trial (NCT06881784) ^5, 6^. Despite these advances, effectiveness of daraxonrasib and other RAS(ON) TCIs has been limited by acquired mechanisms of resistance ^2, 7–11^. In patients with RAS mutant cancers, resistance to daraxonrasib has been associated with acquired mutations that (i) disrupt daraxonrasib binding to RAS, or (ii) enhance native RAS-RAF signaling ^12^. Otherwise, less is known about non-genetic mechanisms of resistance to RAS(ON) TCIs.

Treatment of patients with NSCLC KRAS(G12C) mutant tumors has been advanced with the allele-selective sotorasib and adagrasib inhibitors ^13–16^. However, tolerance to these agents invariably emerges by mechanisms attributed to genomic alterations that converge on reactivation of the MAPK pathway ^7, 8, 10, 17, 18^. Interestingly, sotorasib/adagrasib resistance is circumvented by daraxonrasib and the related RMC-7977 RAS(ON) TCI ^1, 2, 8, 19^. These findings have been attributed to daraxonrasib-mediated inhibition of RAS reactivation in resistance to KRAS(G12C) inhibitors. Whether resistance to sotorasib and daraxonrasib is conferred by complementary mechanisms is not known.

The *MUC1* gene encodes a transmembrane oncogenic MUC1-C/M1C protein that drives resistance of NSCLC KRAS(G12C) mutant cells to sotorasib and adagrasib ^20^. MUC1 evolved to protect barrier tissue cells, such as those lining the respiratory tract, from loss of homeostasis ^21–24^. MUC1 is activated by biotic and abiotic insults with the induction of inflammatory, proliferative and epigenetic pathways that are adaptive responses in wound healing ^21–24^. As a maladaptation of this protective function, prolonged M1C activation in settings of chronic inflammation establishes heritable epigenetic reprogramming that contribute to the NSCLC stem cell (CSC) state and drug resistance ^21–27^. M1C promotes chronic inflammation by binding directly to STAT1 and activating expression of type I interferon (IFN)-stimulated genes (ISGs) ^22–24^. In NSCLC KRAS(G12C) mutant cells, M1C exploits activation of this STAT1-dependent inflammatory response in driving sotorasib/adagrasib resistance ^20^.

There is no reported evidence that M1C plays a role in tolerance to daraxonrasib. The present work in NSCLC KRAS(G12C) cell line and patient-derived models demonstrates that M1C is necessary for daraxonrasib resistance. We report that M1C confers resistance to daraxonrasib by forming cell membrane biocondensates with SHP2 that integrate activation of the downstream OSM/gp130/STAT3 and NF-κB/EMT pathways. Our results reveal that M1C induces resistance to sotorasib/adagrasib and daraxonrasib by noncongruent inflammatory mechanisms. These findings identify M1C as a potential target for treatment of patients with NSCLC KRAS(G12C) tumors refractory to daraxonrasib.

## Results

### Treatment of NSCLC cells with RAS(ON) TCIs induces M1C expression

In investigating potential involvement of M1C in response to RAS(ON) TCIs, we found that treatment of NSCLC H358 KRAS(G12C) cells with daraxonrasib induces M1C mRNA (Fig. 1A; Supplemental Fig. S1A) and protein levels (Fig. 1B). NSCLC H2122 KRAS(G12C) (Fig. 1C; Supplemental Fig. S1B) and patient-derived MGH1112 KRAS(G12C) (Supplemental Fig. S1C) cells also responded to daraxonrasib with induction of M1C expression. Similar results were obtained with the related RMC-7977 inhibitor (Supplemental Figs. S1D and S1E), demonstrating that M1C is induced by different RAS(ON) TCIs.

**Figure 1.**
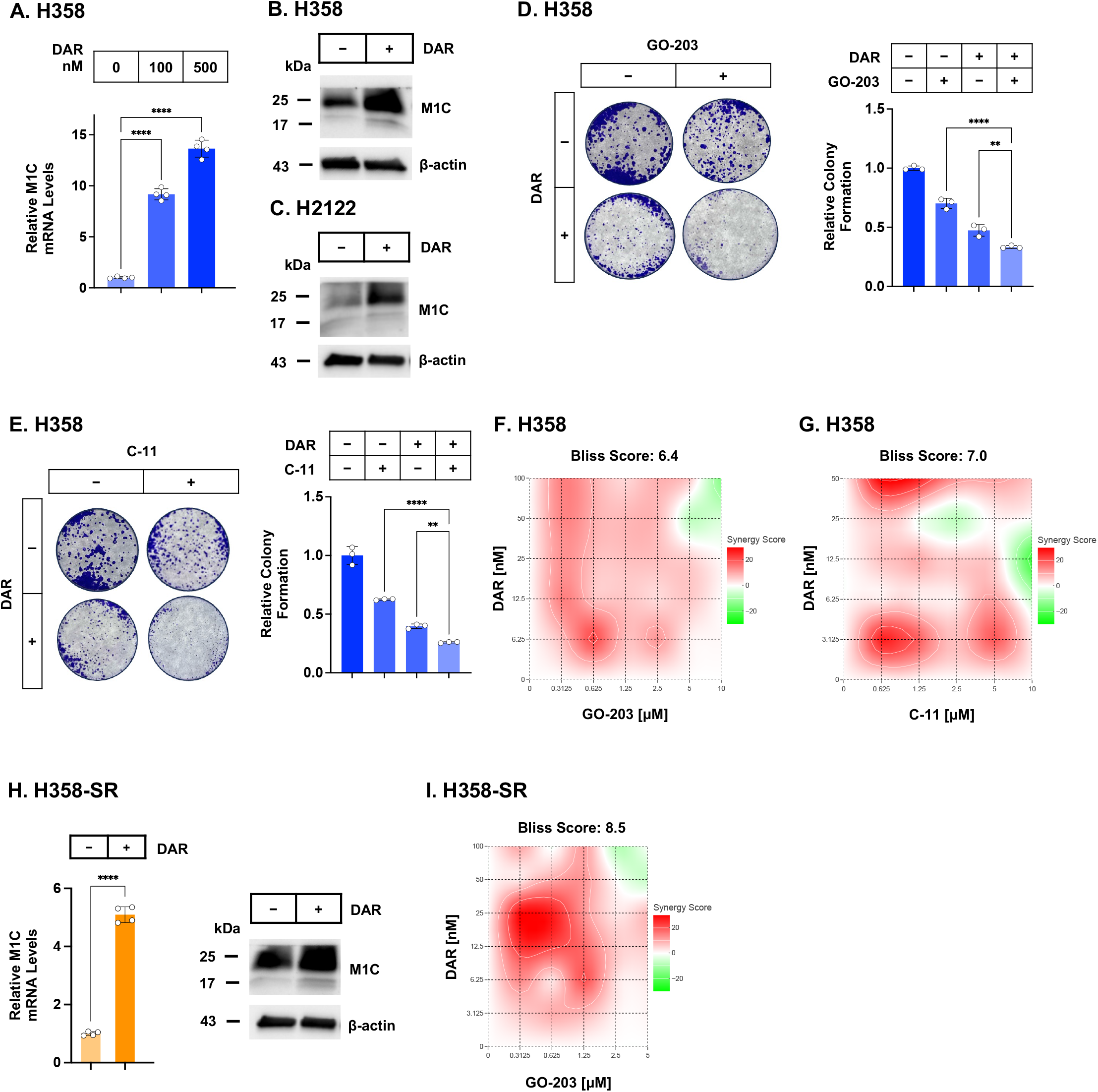
M1C expression is induced by daraxonrasib to mitigate loss of survival. **A**. H358 KRAS G12C cells treated with the indicated concentrations of daraxonrasib for 48 hours were analyzed for M1C transcripts. The results (mean±SD of 4 determinations) are expressed as relative levels compared to that obtained for control cells (assigned a value of 1). **B and C**. Lysates from H358 (**B**) and H2122 (**C**) cells treated with 100 nM daraxonrasib for 48 hours were immunoblotted with antibodies against the indicated proteins. **D**. H358 cells treated with 100 nM daraxonrasib alone, 5 μM GO-203 alone and the combination of these agents for 48 hours were analyzed for colony formation. Shown are representative photomicrographs of stained colonies. The results (mean SD of three determinations) are expressed as relative colony formation compared to that for control cells (assigned a value of 1). **E**. H358 cells treated with 100 nM daraxonrasib alone, 5 μM C-11 alone and the combination of these agents for 48 hours were analyzed for colony formation. Shown are representative photomicrographs of stained colonies. The results (mean±SD of three determinations) are expressed as relative colony formation compared to that for control cells (assigned a value of 1). **F and G**. H358 cells treated with the indicated concentrations of daraxonrasib and GO-203 (**F**) or daraxonrasib and C-11 (**G**) for 48 hours were analyzed for viability by Alamar blue staining. Indicated are the combination indices determined using Bliss scores. **H**. Sotorasib-resistant H358-SR cells treated with 100 nM daraxonrasib for 48 hours were analyzed for M1C transcripts (left). The results (mean±SD of 4 determinations) are expressed as relative levels compared to that obtained for control cells (assigned a value of 1). Lysates were immunoblotted with antibodies against the indicated proteins (right). **I**. H358-SR cells treated with the indicated concentrations of daraxonrasib and GO-203 for 48 hours were analyzed for viability by Alamar blue staining. Indicated are the combination indices determined using Bliss scores.

M1C includes a CQC motif in the cytoplasmic domain that is necessary for M1C homodimerization and function ^28^. Sensitivity of H358 cells to daraxonrasib was increased by targeting the M1C CQC motif with the cell penetrating GO-203 peptide (Fig. 1D) ^23^ and C-11 small molecule (Fig. 1E) ^29^ inhibitors. Combining daraxonrasib with GO-203 (Fig. 1F) and C-11 (Fig. 1G) exhibited synergistic activity as determined by BLISS scores. Corroborating results were obtained with RMC-7977 (Supplemental Figs. S1F and S1G).

Sotorasib-resistant NSCLC KRAS G12C cells retain sensitivity to RAS(ON) TCIs ^1, 2, 8, 19^. We found that sotorasib-resistant H358-SR cells ^20^ respond to daraxonrasib with induction of M1C expression (Fig. 1H). M1C was also induced by daraxonrasib treatment of (i) H2122-SR cells (Supplemental Fig. S1H), and (iii) patient-derived MGH1112 cells intrinsically resistant to sotorasib (Supplemental Fig. S1C). Targeting M1C in H358-SR cells synergistically increased sensitivity to daraxonrasib treatment (Fig. 1I; Supplemental Fig. S1I), indicating M1C is induced as a protective response against RAS(ON) TCIs.

### M1C is necessary for acquired RAS(ON) TCI resistance

To determine if M1C plays a role in daraxonrasib resistance, we selected H358 cells for growth in the presence of increasing daraxonrasib concentrations over 12 months (H358, IC50=19 nM; H358/DAR-R, IC50=593 nM) (Supplemental Fig. S2A). Silencing M1C in H358/DAR-R cells (Supplemental Fig. S2B) enhanced daraxonrasib activity as assessed by inhibition of clonogenicity (Fig. 2A). For comparison, H358 cells were selected for resistance to RMC-7977 (H358/RMC-7977-R, IC50=1188; Supplemental Fig. S2C). Silencing M1C in H358/RMC-7977-R cells (Supplemental Fig. S2D) similarly enhanced sensitivity to RMC-7977-induced loss of colony formation (Supplemental Fig. S2E).

**Figure 2.**
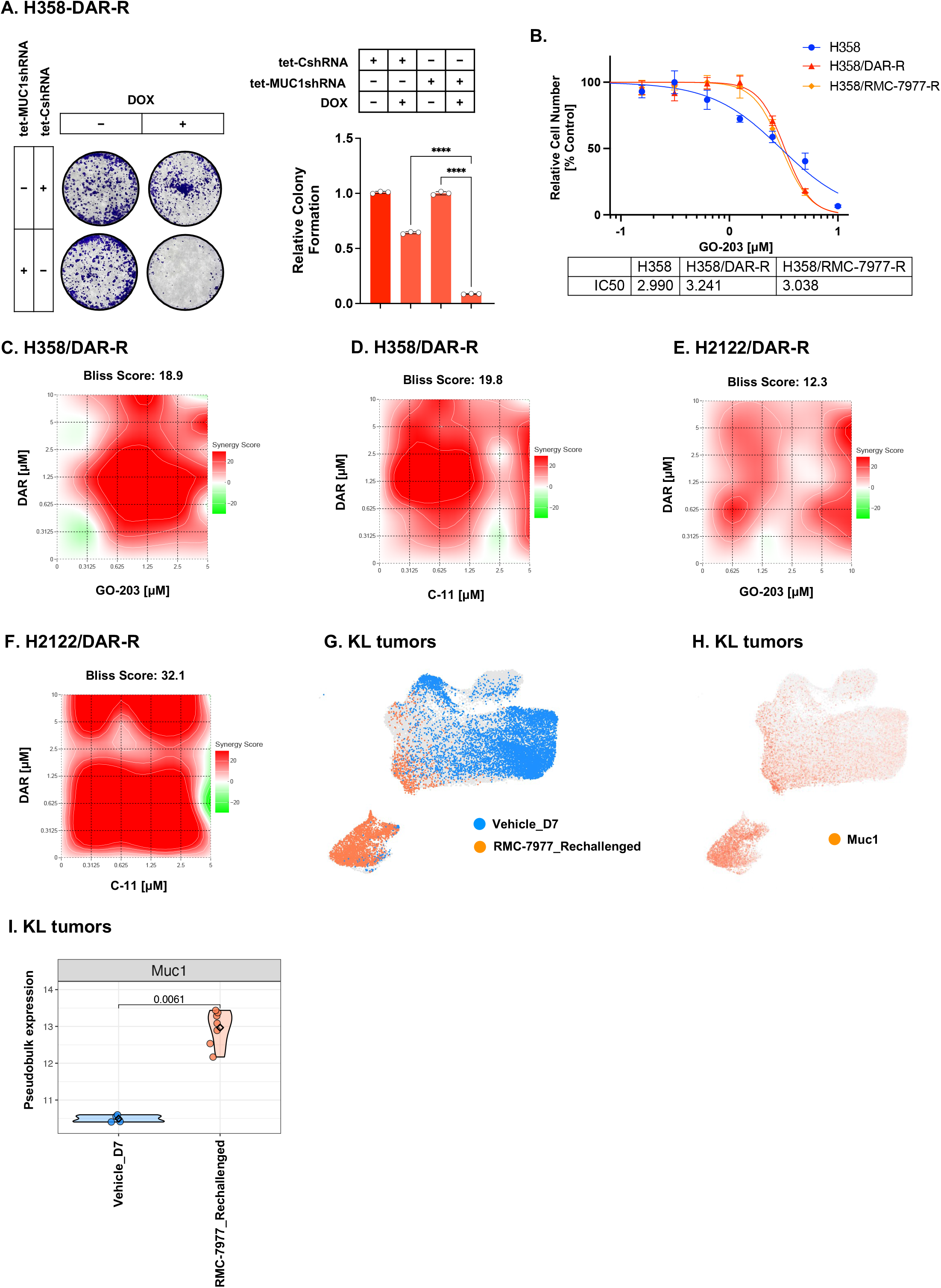
M1C is required for acquired daraxonrasib resistance. **A**. H358/DAR-R/tet-CshRNA and H358/DAR-R/tet-MUC1shRNA cells were treated with vehicle or DOX for 7 days. The cells were then incubated with 100 nM daraxonrasib for 2 days and analyzed for colony formation. Shown are representative photomicrographs of stained colonies. The results (mean±SD of three determinations) are expressed as relative colony number compared to that for control cells (assigned a value of 1). **B**. H358, H358/DAR-R and H358/RMC-7977-R cells treated with the indicated concentrations of GO-203 for 48 hours were analyzed for viability by Alamar blue staining. Indicated are the IC50 values. **C and D**. H358/DAR-R cells treated with the indicated concentrations of daraxonrasib and GO-203 (**C**) or daraxonrasib and C-11 (**D**) for 48 hours were analyzed for viability by Alamar blue staining. Indicated are the combination indices as determined using Bliss scores. **E and F**. H2122/DAR-R cells treated with the indicated concentrations of daraxonrasib and GO-203 (**E**) or daraxonrasib and C-11 (**F**) for 48 hours were analyzed for viability by Alamar blue staining. Indicated are the combination indices determined using Bliss scores. **G**. UMAP of scRNA-seq data from KL tumors (i) treated with vehicle (blue) and (ii) rechallenged with RMC-7977 after drug treatment, withdrawal and disease recurrence (orange). **H**. UMAP of scRNA-seq data with normalized Muc1 expression. **I**. Pseudobulk expression analysis of Muc1 in KL tumor cells treated with vehicle and rechallenged with RMC-7977.

Targeting M1C with GO-203 demonstrated that H358/DAR-R and H358/RMC-7977-R cells are dependent on M1C for survival (Fig. 2B). In addition, we found that treatment of H358/DAR-R cells with GO-203 in combination with daraxonrasib is synergistic as evidence of reversing the daraxonrasib-resistant phenotype (Fig. 2C). Treatment of H358/DAR-R cells with C-11 also reversed daraxonrasib resistance (Fig. 2D).

Consistently, GO-203 and C-11 reversed resistance of H358/RMC-7977-R cells to RMC-7977 (Supplemental Figs. S2F and S2G). To extend these results, we selected H2122 KRAS(G12C) cells for resistance to daraxonrasib (H2122, IC50=1 nM; H2122/DAR-R, IC50=1520 nM) (Supplemental Fig. S2H) and found that treatment with GO-203 (Fig. 2E) and C-11 (Fig. 2F) reverses the H2122/DAR-R daraxonrasib-tolerant cell phenotype.

As an additional model, patient-derived NSCLC MGH1112 KRAS(G12C) cells, which are tolerant to sotorasib (IC50>1 μM)^30^, were selected for resistance to daraxonrasib (MGH1112 cells, IC50=0.5 μM; MGH1112/DAR-R, IC50=88 μM)(Supplemental Fig. S2I). Here again, targeting M1C in MGH1112/DAR-R cells with GO-203 (Supplemental Fig. S2J) and C-11 (Supplemental Fig. S2K) was synergistic with daraxonrasib in reversing resistance.

To extend these findings in an *in vivo* model, we analyzed scRNA-seq datasets obtained from mouse NSCLC KL2 (Kras G12C, STK11-deficient) tumors treated with RMC-7977 ^8^. Treatment with RMC-7977 resulted in tumor regressions that were maintained with continuous exposure for 60 days ^8^. Upon RMC-7977 withdrawal on day 60 and subsequent rechallenge, we found that the persisting tumor cells are enriched for induction of Muc1 expression (Figs. 2G and 2H). These results were extended by pseudobulk expression analysis, which confirmed upregulation of Muc1 transcripts in the KL2 long-term persister tumor cell population (Fig. 2I).

These findings collectively indicate that M1C promotes resistance of NSCLC KRAS(G12C) cells to the daraxonrasib and RMC-7977 RAS(ON) TCIs.

### M1C forms cell membrane condensates in daraxonrasib-tolerant cells

Membraneless intracellular biomolecular condensates are formed by liquid-liquid phase separation (LLPS) in cancer cells ^31^. M1C (i) is modified by galectin-3, (ii) includes a cytoplasmic intrinsically disordered region (IDR), (iii) forms multimers and (iv) binds to RNPs, which are characteristics that collectively contribute to condensate formation ^23, 24, 32^. To our knowledge, there is no reported association of LLPS structures with daraxonrasib resistance. We identified M1C punctate structures in H358/DAR-R, as compared to parental H358, cells (Fig. 3A). Similar M1C puncta were detected in H2122/DAR-R cells (Supplemental Fig. S3A). Treatment with 1,6-hexanediol abolished these puncta (Supplemental Fig. S3B), consistent with biomolecular condensates ^33^. Notably, targeting M1C with GO-203 and C-11 disrupted these puncta (Fig. 3B), indicating that M1C is necessary for their formation. Analysis of H358/DAR-R cells further demonstrated that M1C-dependent puncta are associated with the cell membrane as determined by imaging at higher magnification (Fig. 3C) and colocalization with the cell membrane Na+/K+ ATPase alpha 1 marker (Supplemental Fig. S3C).

**Figure 3.**
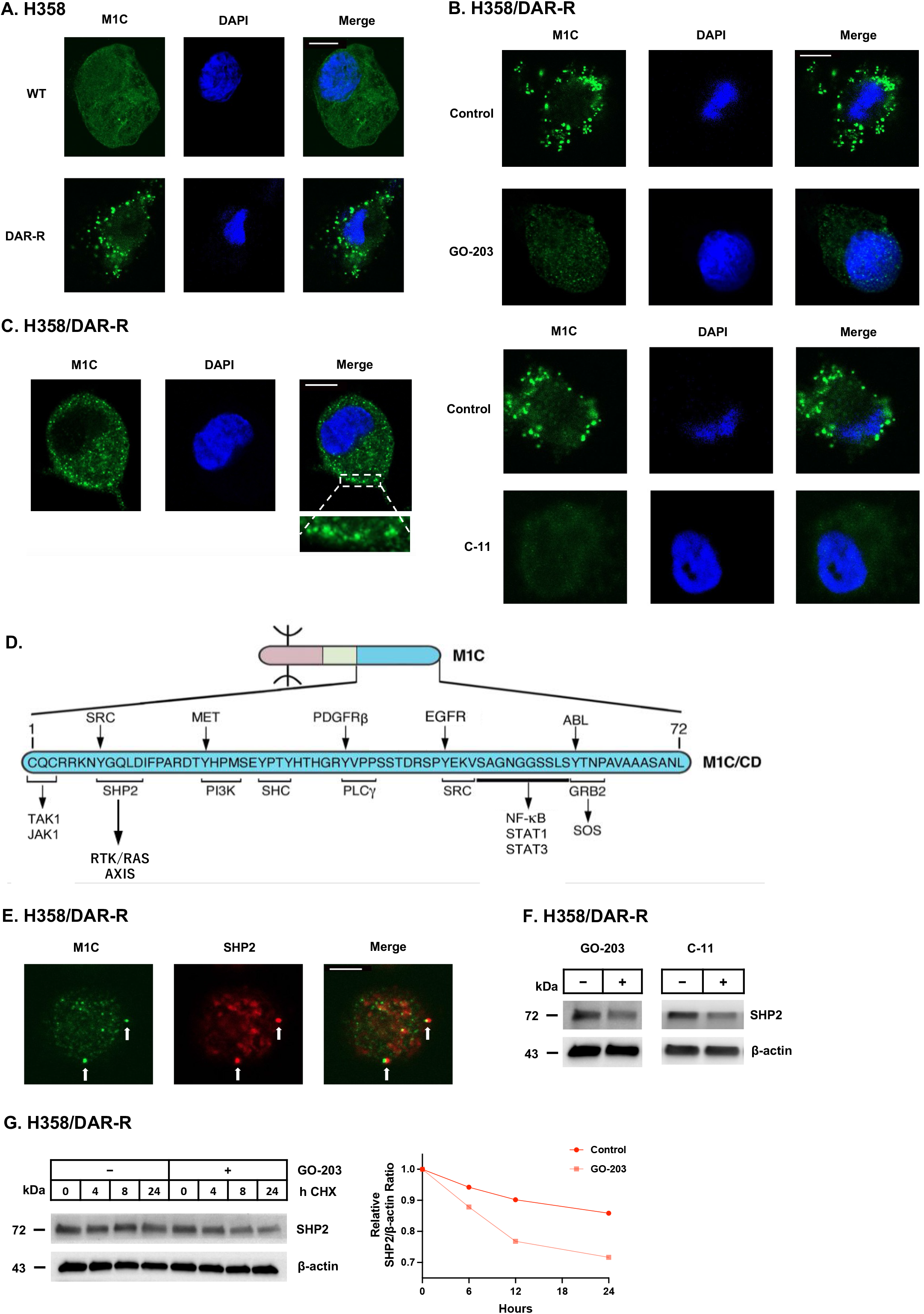
M1C forms biomolecular condensates with SHP2 in daraxonrasib-resistant cells. **A**. Confocal IF imaging of H358 and H358/DAR-R cells showing localization of M1C in extra-nuclear puncta in association with daraxonrasib resistance. Nuclei were stained with DAPI. Scale bar, 5 µm. **B**. Confocal IF imaging of H358/DAR-R cells left untreated and treated with 5 μM GO-203 or 5 μM C-11 for 48 hours showing effects on M1C localization in puncta. Scale bar, 5 µm. **C**. Confocal IF imaging of H358/DAR-R cells showing localization of M1C puncta at the cell membrane. Scale bar, 5 µm. **D**. Schema of the M1C IDR with highlighting of TK phosphorylation sites and binding of effectors that regulate RTK/RAS signaling. The M1C IDR also has the capacity for mutually exclusive activation of the inflammatory JAK®STAT1/3 and TAK1®NF-κB pathways. **E**. Confocal IF imaging of H358/DAR-R cells showing co-localization of M1C and SHP2 in condensates. Scale bar, 5 µm. **F**. Lysates from H358/DAR-R cells treated with 5 μM GO-203 or 5 μM C-11 for 48 hours were immunoblotted with antibodies against the indicated proteins. **G**. H358/DAR-R cells were treated with cycloheximide alone or in combination with 5 μM GO-203 for the indicated hours. Lysates were immunoblotted with antibodies against the indicated proteins. Intensity of the IB bands determined by densitometry is expressed as the relative SHP2/β-actin ratio.

The M1C cytoplasmic protein is a 72 aa IDR that acts as a scaffold for effectors of the RTK/RAS axis (Fig. 3D) ^23, 24^. The M1C 72 aa IDR forms a complex with the SHP2 protein tyrosine phosphatase ^34^, which is required to fully activate the RTK/RAS/ERK axis (Fig. 3D) ^35^. Phosphorylation of the YGQLD motif on Tyr functions as a binding site for the SHP2 N-terminal SH2 domain in activating RTK-mediated RAS signaling (Fig. 3D) ^34^. Noteworthy is that SHP2 also contributes to LLPS through interactions at its PTP domain ^36, 37^.

Consistent with the direct interaction between M1C and SHP2 ^34^, we found that M1C colocalizes with SHP2 in H358/DAR-R cell puncta (Fig. 3E). Targeting M1C in H358/DAR-R cells suppressed SHP2 protein levels in the absence of decreases in SHP2 mRNA transcripts (Fig. 3F; Supplemental Fig. S3D). In support of a post-transcriptional mechanism, we found that M1C stabilizes the SHP2 protein (Fig. 3G). We also found that treatment with the allosteric SHP2 inhibitors TNO155 (batoprotafib) ^38^(Supplemental Fig. S3E) and SHP099 ^39, 40^ (Supplemental Fig. S3F) reverses daraxonrasib resistance, indicating that M1C/SHP2 signaling drives the daraxonrasib-resistant phenotype.

### M1C activates the STAT3 pathway in conferring daraxonrasib tolerance

M1C-driven resistance of NSCLC KRAS G12C cells to sotorasib/adagrasib is dependent on induction of the inflammatory STAT1 pathway ^20^. The M1C IDR interacts with (i) JAK1 at a site adjacent to that for SHP2 binding and (ii) STAT1/3 at a downstream region (Fig. 3D) in acting as a scaffold facilitating JAK1-STAT1/3 activation ^23, 24^. There is no known involvement of M1C in integrating the SHP2 and STAT pathways.

Analysis of RNA-seq datasets from H358/DAR-R vs H358 cells demonstrated downregulation of the STAT1-driven HALLMARK INTERFERON (IFN) ALPHA and GAMMA RESPONSE signatures (Supplemental Fig. S4A). In contrast to activation of the STAT1 transcriptome in sotorasib/adagrasib resistance ^20^, daraxonrasib tolerance was associated with upregulation of the HALLMARK IL6 JAK STAT3 SIGNALING gene signature (Fig. 4A). Comparison of H358/DAR-R vs H358 cells further identified decreases in STAT1 and increases in STAT3 expression (Fig. 4B). M1C binds directly to STAT3 and regulates STAT3 target genes that include activation of *MUC1* in an auto-inductive pathway ^41^. In concert with involvement of the M1C/STAT3 pathway, targeting M1C in H358/DAR-R cells with GO-203 and C-11 downregulated STAT3 levels (Supplemental Fig. S4B).

**Figure 4.**
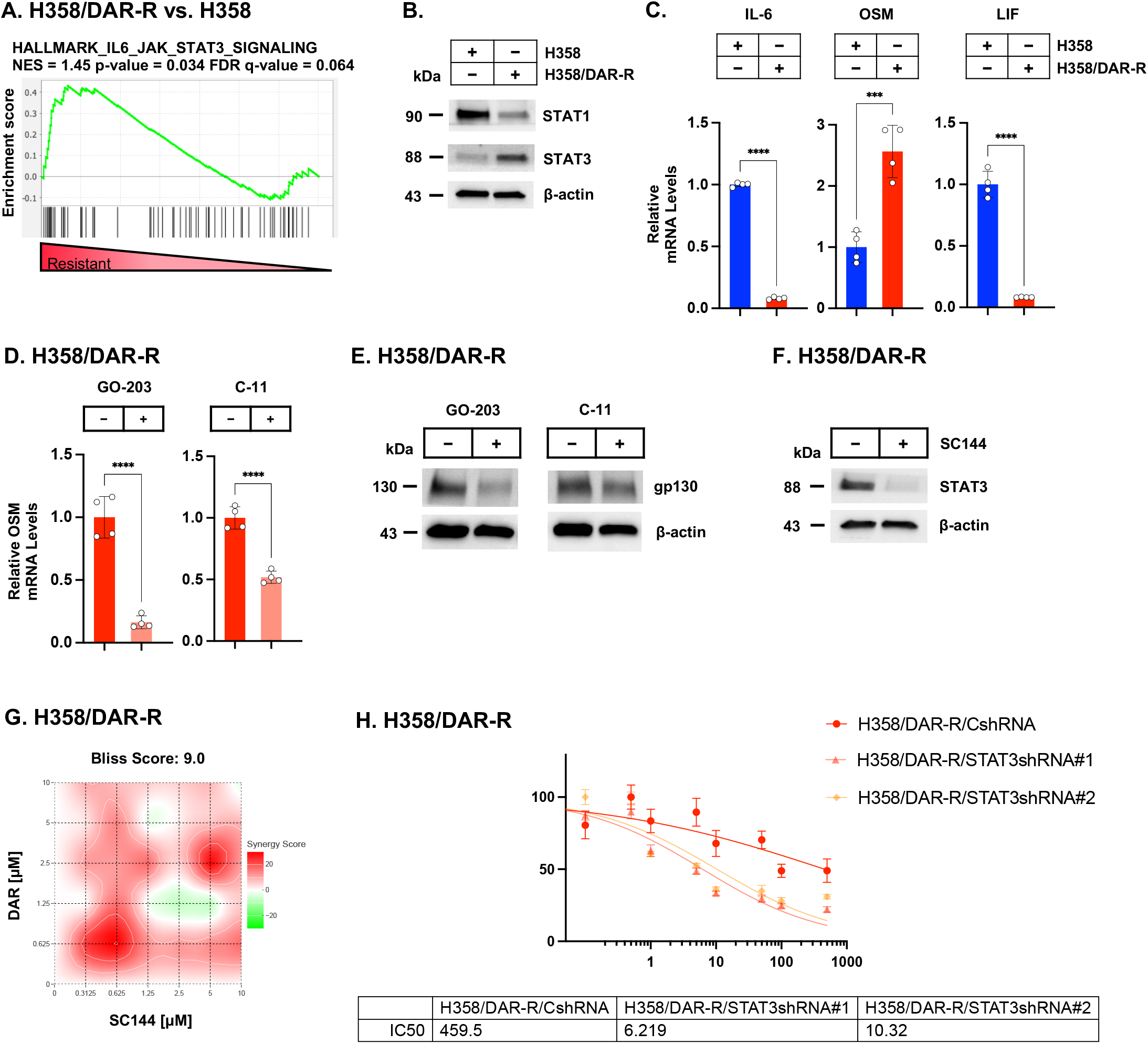
M1C activates the OSM/gp-130/STAT3 pathway in driving daraxonrasib resistance. **A**. RNA-seq datasets from H358/DAR-R vs H358 cells were analyzed using the HALLMARK IL6 JAK STAT3 SIGNALING gene signature. **B**. Lysates from H358 and H358/DAR-R cells were immunoblotted with antibodies against the indicated proteins. **C**. H358 and H358/DAR-R cells were analyzed for IL-6, OSM and LIF transcripts. The results (mean±SD of 4 determinations) are expressed as relative levels compared to that obtained for H358 cells (assigned a value of 1). **D**. H358/DAR-R cells (i) treated with 5 μM GO-203 or 5 μM C-11 for 48 hours were analyzed for OSM transcripts. The results (mean±SD of 4 determinations) are expressed as relative levels compared to that obtained for control cells (assigned a value of 1). **E**. Lysates from H358/DAR-R cells treated with 5 μM GO-203 or 5 μM C-11 for 48 hours were immunoblotted with antibodies against the indicated proteins. **F**. Lysates from H358/DAR-R cells treated with 5 μM SC144 for 48 hours were immunoblotted with antibodies against the indicated proteins. **G**. H358/DAR-R cells treated with the indicated concentrations of daraxonrasib and SC144 for 48 hours were analyzed for viability by Alamar blue staining. Indicated are the combination indices determined using Bliss scores. **H**. H358/DAR-R cells stably expressing a CshRNA, STAT3shRNA#1 or STAT3shRNA#2 and treated with the indicated concentrations of daraxonrasib for 48 hours were analyzed for viability by Alamar blue staining. Indicated are the IC50 values.

STAT3 is activated by the IL-6 family of cytokines that include IL-6, oncostatin m (OSM) and leukemia inhibitory factor (LIF) ^42^. We found that daraxonrasib resistance is associated with (i) marked upregulation of OSM, and (ii) downregulation of IL-6 and LIF expression (Fig. 4C). The OSM cytokine (i) activates MAPK signaling in contrast to IL-6 and LIF, and (ii) induces separable stemness and mesenchymal programs in cancer cells ^43^. We therefore focused on the OSM signaling pathway, which is activated by binding of gp130 to the OSMR and LIFR receptors ^42^. Like OSM, gp130 expression was increased in H358/DAR-R vs H358 cells (Supplemental Fig. S4C). Targeting M1C in H358/DAR-R cells suppressed OSM (Fig. 4D) and gp-130 (Fig. 4E) expression, indicating that M1C induces OSM, gp130 and STAT3.

In investigating whether the M1C-driven OSM/gp130/STAT3 pathway confers tolerance to daraxonrasib, we found that treatment of H358/DAR-R cells with the gp130 inhibitor SC144 ^44^ downregulates STAT3 (Fig. 4F) and reverses daraxonrasib resistance (Fig. 4G). Similar results were observed with H2122/DAR-R (Supplemental Fig. S4D) cells, indicating that the OSM/gp130/STAT3 pathway contributes to the daraxonrasib-resistant phenotype. By extension, silencing STAT3 in H358/DAR-R cells (Supplemental Fig. S4E) reversed daraxonrasib resistance (Fig. 4H). We also found that (i) inhibiting gp130 in H358/DAR-R cells with SC144 suppresses SHP2 mRNA and protein levels (Supplemental Figs. S4F and S4G), and (ii) silencing STAT3 downregulates SHP2 expression (Supplemental Fig. S4H), indicating that M1C integrates activation of the SHP2 and OSM/gp130/STAT3 pathways in association with daraxonrasib resistance.

### M1C induces EMT in driving daraxonrasib tolerance

M1C and SHP2 each promote EMT in NSCLC cells ^25, 45, 46^. Analysis of the H358/DAR-R vs H358 cell transcriptomes identified induction of the HALLMARK EMT gene signature (Fig. 5A). Compared to H358 cells and in concert with an EMT phenotype, H358/DAR-R cells exhibited (i) a spindle-like morphology (Fig. 5B), and (ii) decreases in E-cadherin and increases in vimentin and ZEB1 expression (Supplemental Fig. S5A).

**Figure 5.**
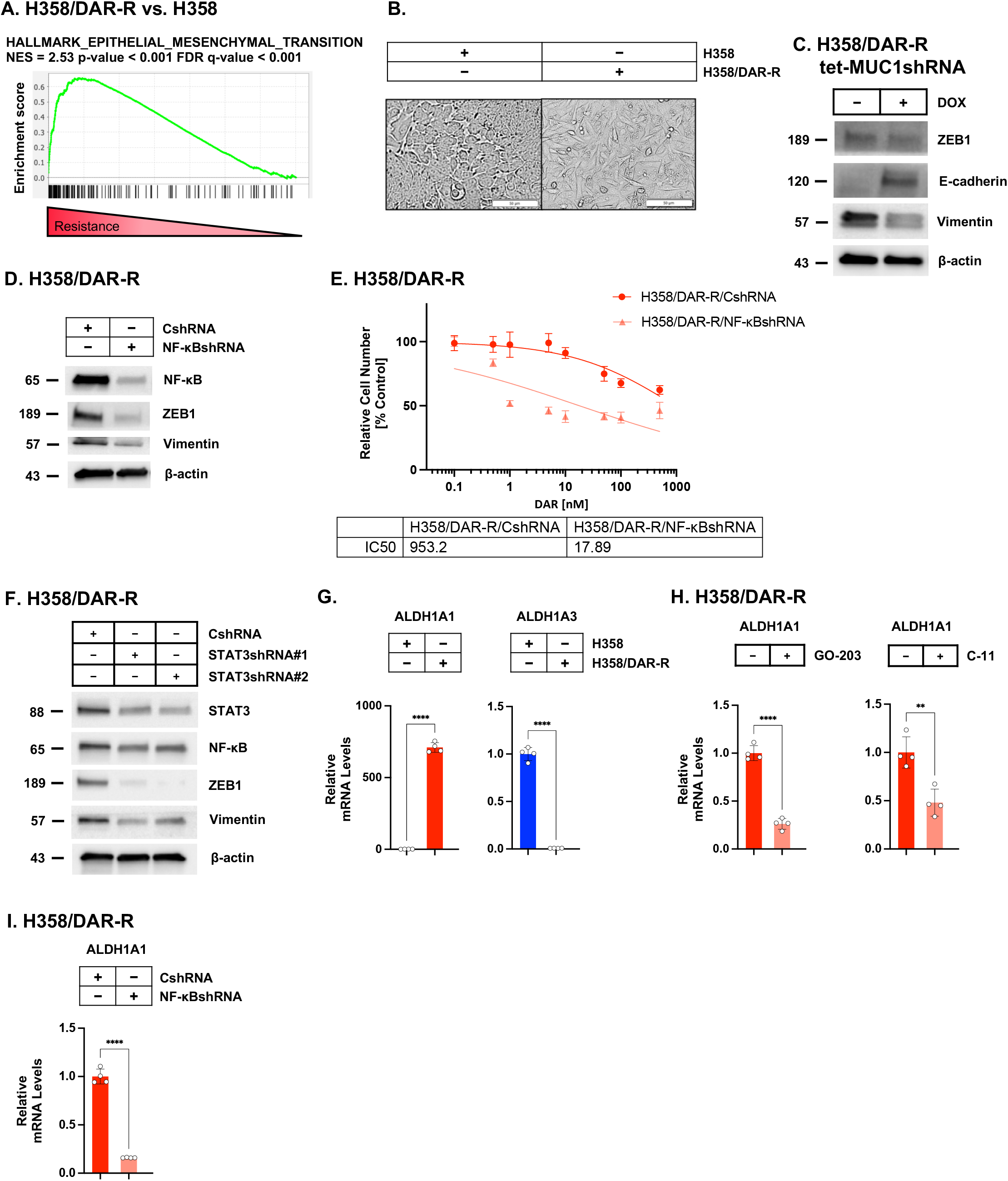
M1C drives EMT by an NF-κB-mediated pathway in conferring daraxonrasib resistance. **A**. Analysis of H358/DAR-R vs H358 cell transcriptomes using the HALLMARK EPITHELIAL MESENCHYMAL TRANSITION gene signature. **B**. Morphology of H358 and H358/DAR-R cells analyzed by microscopy. **C**. Lysates from H358/DAR-R/tet-MUC1shRNA cells expressing treated with vehicle or DOX for 7 days were immunoblotted with antibodies against the indicated proteins. **D**. Lysates from H358/DAR-R/CshRNA or H358/DAR-R/NF-κBshRNA cells were immunoblotted with antibodies against the indicated proteins. **E**. H358/DAR-R/CshRNA and H358/DAR-R/NF-κBshRNA cells treated with the indicated concentrations of daraxonrasib for 48 hours were analyzed for viability by Alamar blue staining. Indicated are the IC50 values. **F**. H358 and H358/DAR-R cells were analyzed for ALDH1A1 and ALDH1A3 transcripts. The results (mean±SD of 4 determinations) are expressed as relative levels compared to that obtained for H358 cells (assigned a value of 1). **G**. H358/DAR-R cells treated with 5 μM GO-203 or 5 μM C-11 for 48 hours were analyzed for ALDH1A1 transcripts. The results (mean±SD of 4 determinations) are expressed as relative levels compared to that obtained for control cells (assigned a value of 1). **G**. H358 and H358/DAR-R cells were analyzed for ALDH1A1 transcripts. The results (mean±SD of 4 determinations) are expressed as relative levels compared to that obtained for H358 cells (assigned a value of 1). **H**. H358/DAR-R/CshRNA and H358/DAR-R/NF-κBshRNA cells were analyzed for ALDH1A1 transcripts (right). The results (mean±SD of 4 determinations) are expressed as relative levels compared to that obtained for CshRNA cells (assigned a value of 1).

The M1C intrinsically disordered region (IDR) interacts directly with TAK1 and NF-kB p65 (RELA) in activating the NF-kB pathway (Fig. 3D)^47, 48^. Within the M1C IDR, (i) TAK1 binds to the same site as JAK1, and (ii) NF-kB interacts with the same motif as STAT3 (Fig. 3D), indicating mutually exclusive activation of these pathways.

M1C regulates NF-kB-target genes that include MUC1 in an auto-inductive loop and ZEB1 in driving EMT ^47, 49^. Silencing M1C in H358/DAR-R cells suppressed (i) ZEB1 expression, and (ii) the EMT phenotype as evidenced by upregulation of E-cadherin and downregulation of vimentin levels (Fig. 5C). Similar results were observed when targeting M1C in H358/DAR-R cells with GO-203 and C-11 (Supplemental Fig. S5B).

Silencing NF-κB in H358/DAR-R cells also suppressed (i) ZEB1 and vimentin levels (Fig. 5D), (ii) the spindle-like morphology (Supplemental Fig. S5C), and (iii) daraxonrasib resistance (Fig. 5E). In extending these results to H2122/DAR-R cells, NF-κB was necessary for (i) ZEB1 expression, (ii) EMT as evidenced by suppression of E-cadherin (Supplemental Fig. S5D), and (iii) daraxonrasib resistance (Supplemental Fig. S5E). As found for NF-κB dependence, STAT3 was also necessary for induction of ZEB1 and EMT (Fig 5F).

ALDH1A1 is a key effector of EMT and drug resistance ^50^. We found that ALDH1A1, but not the related ALDH1A3 isoform, is markedly upregulated in H358/DAR-R vs H358 cells (Fig. 5G). ALDH1A1 expression was also increased in (i) H2122/DAR-R vs H2122 cells (Supplemental Fig. S5F) and (ii) MGH1112/DAR-R vs MGH1112 cells (Supplemental Fig. S5G). Targeting M1C suppressed ALDH1A1 expression in H358/DAR-R cells (Fig. 5H). Moreover, we found that NF-κB is necessary for the upregulation of ALDH1A1 in H358/DAR-R cells (Fig. 5I). These findings demonstrate that M1C integrates NF-κB- and STAT3-mediated induction of EMT in daraxonrasib resistance.

### M1C is a target for elimination of NSCLC CSCs tolerant to daraxonrasib

EMT is intricately interconnected with the CSC state and self-renewal ^51^. In determining if M1C integrates daraxonrasib tolerance with the capacity for self-renewal, we found that targeting M1C in H358/DAR-R cells with GO-203 (Fig. 6A) and C-11 (Fig. 6B) suppresses tumorsphere formation. Comparable results were obtained with H2122/DAR-R (Fig. 6C; Supplemental Fig. S6A) and MGH1112/DAR-R (Fig. 6D; Supplemental Fig. S6B) cells, indicating that M1C is necessary for self-renewal capacity of daraxonrasib-resistant cells.

**Figure 6.**
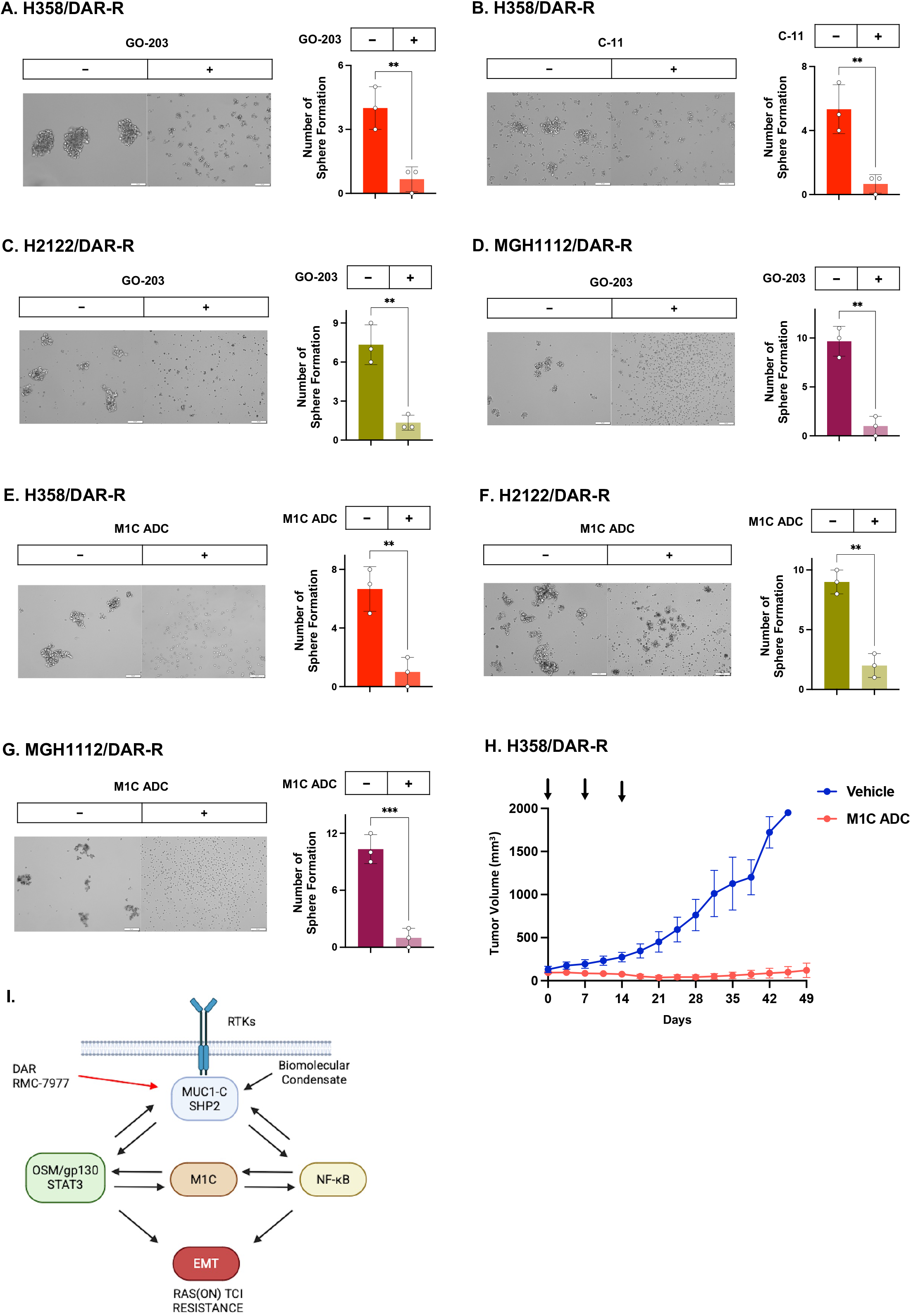
M1C is necessary for self-renewal capacity and tumorigenicity of daraxonrasib-resistant cells. **A and B**. H358/DAR-R cells treated with 5 μM GO-203 (**A**) or 5 μM C-11 (**B**) for 48 hours were analyzed for tumorsphere formation. Shown are representative photomicrographs of tumorspheres (left). The results (mean±SD of three determinations) are expressed as relative tumorsphere formation compared to that for vehicle treated cells (assigned a value of 1) (right). **C and D**. H2122/DAR-R (**C**) and MGH1112/DAR-R (**D**) cells treated with 5 μM GO-203 for 48 hours were analyzed for tumorsphere formation. Shown are representative photomicrographs of tumorspheres (left). The results (mean±SD of three determinations) are expressed as relative tumorsphere formation compared to that for vehicle treated cells (assigned a value of 1) (right). **E-G**. H358/DAR-R (**E**), H2122/DAR-R (**F**) and MGH1112/DAR-R (**G**) cells treated with vehicle or 50 nM M1C ADC for 7 days were analyzed for tumorsphere formation. Shown are representative photomicrographs of tumorspheres (left). The results (mean±SD of three determinations) are expressed as number of tumorsphere formations compared to that for vehicle treated cells (assigned a value of 1) (right). **H**. Nude mice were injected subcutaneously with 5 × 10^6^ H358/DAR-R cells. Mice were randomized into two groups when the mean tumor volume reached 100 mm^3^ and then treated with vehicle (n=5) or 7.5 mg/kg M1C ADC weekly x 3 (n=5). Tumor volumes are expressed as the mean±SEM. **I**. Proposed model depicting M1C-induced signaling in daraxonrasib resistance. M1C forms cell membrane-associated condensates with SHP2 in facilitating SHP2 stabilization. M1C/SHP2 signaling activates the (i) OSM/gp130/STAT3 and (ii) NF-κB p65 pathways. Binding of M1C directly to STAT3 and NF-κB regulates activation of their target genes which include *MUC1* in auto-inductive inflammatory pathways. In turn, STAT3 and NF-κB/ZEB1 drive EMT and RAS(ON) TCI resistance.

Given these results, we asked if M1C is a potential target for eliminating NSCLC CSCs refractory to daraxonrasib. Along these lines, a M1C ADC has been generated with an antibody against the M1C extracellular domain conjugated to the MMAE payload ^52^. Consistent with upregulation of M1C in association with daraxonrasib resistance, we found that sensitivity to the M1C ADC is increased against H358/DAR-R cells as compared to parental H358 cells (Supplemental Fig. S6C). The M1C ADC was also effective in inhibiting self-renewal capacity of H358/DAR-R (Fig. 6E), H2122/DAR-R (Fig. 6F) and MGH1112/DAR-R (Fig. 6G) cells. Moreover, treatment with the M1C ADC suppressed growth of H358/DAR-R tumor xenografts (Fig. 6H) in the absence of significant weight loss and other overt toxicities (Supplemental Fig. S6D). These findings demonstrate that M1C is necessary for self-renewal and tumorigenicity of daraxonrasib-tolerant NSCLC KRAS(G12C) cells.

## Discussion

The selective KRAS(G12C) sotorasib/adarasib inhibitors have substantially advanced NSCLC treatment ^53^. However, resistance due to mechanisms that include reactivation of RAS/MAPK signaling by other RAS members have limited their longer-term efficacy ^17, 54, 55^. RAS(ON) TCIs counteract reactivation RAS/MAPK signaling and are effective in settings of sotorasib resistance ^17, 53–55^. As found for the KRAS(G12C)-selective inhibitors, development of resistance to RAS(ON) TCIs has also challenged their effectiveness ^53^. The present studies demonstrate that M1C is induced by treatment of NSCLC KRAS(G12C) cells with daraxonrasib and RMC-7977. Our results further demonstrate that, as uncovered for sotorasib/adagrasib ^20^, M1C is necessary for resistance to daraxonrasib and RMC-7977. Together, these findings demonstrate that M1C confers tolerance to both KRAS(G12C)-selective and RAS(ON) TCIs. Nonetheless and of potential therapeutic relevance, it was not clear if M1C drives resistance to these agents with distinct structures and targets by the same mechanisms.

Our studies reveal that M1C forms biomolecular condensates in NSCLC cells with acquired resistance to daraxonrasib (Fig. 6I). The M1C IDR functions as a scaffold modified by RTKs with the capacity to facilitate activation of effectors, such as SHP2, in the RTK/RAS axis ^23, 24^. We found that M1C forms cell membrane-associated condensates with SHP2 in the setting of daraxonrasib tolerance (Fig. 6I). This observation indicated that maintaining M1C and SHP2 in proximity could contribute to acquisition of the resistant phenotype. In support of this reasoning, M1C stabilized SHP2 and, like M1C, SHP2 was necessary for daraxonrasib resistance. The formation of condensates at the cell membrane has been linked to TKI inhibitor resistance as a mechanism to enhance adaptive RTK signaling ^56^. Given that RAS(ON) TCIs disrupt the RTK/RAS axis, it is conceivable that the condensates identified here are similar in structure and function. In this regard, studies will be needed to determine if M1C-induced resistance of (i) NSCLC KRAS(G12C) cells to sotorasib/adagrasib, and (ii) NSCLC EGFR mutant cells to osimertinib is associated with the formation of M1C-dependent cell membrane condensates ^20, 57^. M1C is necessary for the formation of nuclear paraspeckles in response to DNA replication stress ^32^. The present work provides the first evidence that M1C forms cell membrane-associated condensates in response to inhibiting RAS signaling. We note that these findings do not exclude the possibility that M1C could also contribute to the formation of stress granules in response to targeted agents ^58^.

M1C-dependent activation of the inflammatory STAT1 pathway confers resistance of NSCLC KRAS G12C cells to sotorasib ^20^. M1C/STAT1 signaling is also responsible for resistance of (i) NSCLC EGFR mutant cells to osimertinib ^52, 57^, and (ii) PDAC KRAS G12D cells resistant to MRTX1133 ^59^. These findings provided support for activation of the M1C/STAT1 pathway as a common mechanism for resistance to agents targeting the RTK/RAS axis. Surprisingly, comparison of H358/DAR-R vs H358-SR cell transcriptomes identified (i) downregulation of the IFN ALPHA/GAMMA (Supplemental Fig. S7A), and (ii) upregulation of the HALLMARK E2F TARGETS and HALLMARK G2M CHECKPOINT (Supplemental Fig. S7B) gene signatures, supporting a transition of inflammatory to proliferative signaling in daraxonrasib resistance. In addition, we uncovered that STAT3, an effector of resistance to targeted agents in oncogene-addicted cells ^60^, is essential for conferring the RAS(ON) TCI-resistant phenotype. The basis for this switch from STAT1 to STAT3 was attributable to activation of the OSM/gp130/STAT3 pathway ^61^. The induction of OSM and gp130 in association with daraxonrasib resistance was M1C- and STAT3-dependent, indicating involvement of the M1C/STAT3 auto-inductive pathway ^41^. The novelty of the present work is that there was no previously known involvement of a M1C-driven OSM/gp130/STAT3 pathway in conferring resistance to a RAS(ON) TCI or other targeted agents.

The interrelationships among EMT, the CSC state and drug resistance are well established ^51, 62, 63^. STAT3 and NF-κB signaling are both linked to inducing EMT and targeted drug resistance ^64, 65^. Like STAT3 ^41^, M1C binds directly to NF-κB and regulates the expression of NF-κB target genes ^47^. Our results demonstrate that M1C integrates the STAT3 and NF-κB pathways in driving EMT, the CSC state and RAS(ON) TCI resistance (Fig. 6I). EMT is associated with induction of the ALDH1A1/3 isoforms in conferring drug resistance. Here, we found that daraxonrasib resistance is associated with induction of ALDH1A1 and, conversely, ALDH1A3 is induced in the sotorasib-resistant phenotype (Supplemental Fig. S7C). Collectively, these results indicate that M1C-driven resistance of NSCLC KRAS(G12C) mutant cells to daraxonrasib and sotorasib is largely conferred by noncongruent pathways (Supplemental Fig. S7D). This conclusion is based on studies of isogenic cells established for resistance to these agents.

Our findings support M1C as a target for reversing daraxonrasib resistance. M1C is druggable with an ADC under development by the NCI NExT Program and small molecules that inhibit the M1C cytoplasmic domain. These anti-M1C agents could be effective alone and in combination with daraxonrasib for the treatment of patients with NSCLC KRAS(G12C) tumors refractory to this agent. Targeting M1C could conceivably also delay acquisition of resistance to RAS(ON) TCIs.

## Materials and Methods

### Cell culture

H358 KRAS G12C mutant, TP53 null (ATCC, Manassas, VA, USA) and H2122 KRAS G12C mutant, TP53 null (ATCC) cells were cultured in RPMI 1640 medium (Corning, NY, USA) supplemented with 10% FBS and 2 mM glutamine. MGH1112 (KRAS G12C, STK11 and KEAP1 mutant) cells were maintained in RPMI1640 medium with 10% FBS as described ^52^. Daraxonrasib- and RMC-7977-resistant cells maintained in the presence of these agents were grown in drug-free medium for 24 h before analysis and for 7 days during inducible M1C silencing. Authentication of the cells was performed by short tandem repeat (STR) analysis every 4 months as described ^52^. Cells were monitored with the MycoAlert Mycoplasma Detection Kit (Lonza, Rockland, ME, USA) every 3 months.

### Quantitative reverse-transcription PCR (qRT-PCR)

Total RNA was extracted using TRIzol reagent (Thermo Fisher Scientific, Waltham, MA, USA) as described ^52^. cDNA synthesis was performed with the High-Capacity RNA-to-cDNA Kit (Applied Biosystems, Grand Island, NY, USA). Quantitative PCR was performed using Power SYBR Green PCR Master Mix (Applied Biosystems) as described ^52^. Primer sequences are listed in Supplementary Table S1.

### Immunoblot analysis

Total cell lysates and chromatin prepared from non-confluent cells were analyzed by immunoblotting with anti-M1C (MA5-11202, 1:50 dilution; Invitrogen, Thermo Fisher Scientific, Waltham, MA, USA, RRID:AB_11000874), anti-β-actin (A5441, 1:2000 dilution; Sigma-Aldrich, Burlington, MA, USA, RRID:AB_476744), anti-Histone H3 (9715, 1:1000 dilution; CST, RRID:AB_331563), anti-STAT1 (RAB01893, 1:500 dilution; CST, RRID:AB_3710444), anti-STAT3 (9139, 1:1000 dilution; CST, RRID:AB_331757), anti-SHP2 (3752, 1:1000 dilution; CST, RRID:AB_2300607), anti-gp130 (3732, 1:1000 dilution; CST, RRID:AB_2125953), anti-ERK (9107, 1:1000 dilution; CST, RRID:AB_10695739), anti-p-ERK (4377, 1:1000 dilution; CST, RRID:AB_331775), anti-NF-κB p65 (8242, 1:1000 dilution; CST, RRID:AB_10859369), anti-ZEB1 (3396, 1:1000 dilution; CST, RRID:AB_1904164), anti-E-cadherin (3195, 1:1000 dilution; CST, RRID:AB_2291471) and anti-vimentin (5741, 1:1000 dilution; CST, RRID: AB_10695459) as described ^52^.

### Confocal immunofluorescence analysis

Cells were fixed in 4% paraformaldehyde (Sigma) at room temperature for 10 min. The samples were incubated with 0.1% Triton X-100 (Sigma) at room temperature for 10 min, blocked with 3% Normal Goat Serum (Gibco), incubated with anti-M1C (16564, 1:500 dilution; CST, RRID:AB_2798765), anti-SHP2 (MA5-17160, 1:200 dilution; Invitrogen, RRID:AB_2538631) and anti-alpha 1 Sodium Potassium ATPase antibody (ab7671, 1:100 dilution; Abcam, RRID:AB_306023) at 4°C overnight and then incubated with goat anti-rabbit IgG H&L labeled with Alexa Fluor 488 (ab150077, 1:500 dilution; Abcam, RRID:AB_2630356), goat anti-mouse IgG H&L labeled with Alexa Fluor 647 (ab150115, 1:500 dilution; Abcam, RRID:AB_2687948) at room temperature for 1 h. Invitrogen™ ProLong™ Diamond Antifade Mountant with DAPI (P36966, Invitrogen) was used for staining of nuclei. The cells were analyzed using a Zeiss 980 Confocal microscope and Fiji/ImageJ (NIH, Bethesda, MD, USA; RRID:SCR_002285).

### Gene silencing

MUC1shRNA (MISSION shRNA TRCN0000122938; Sigma) or a control scrambled shRNA (CshRNA; Sigma) was inserted into the pLKO-tet-puro vector (Plasmid #21915; Addgene, Cambridge, MA, USA). STAT3shRNA#1 (TRCN0000329888), STAT3shRNA#2 (TRCN0000353630), NF-κBshRNA#1 (TRCN0000014686) were produced in HEK293T cells as described ^52^. Cells transduced with the vectors were selected for growth in 1-4 mg/ml puromycin as described ^52^. Cells were treated with 0.1% DMSO as the vehicle control or 500 ng/ml doxycycline (DOX; Millipore Sigma).

### RNA-seq analysis

Total RNA from cells cultured separately in triplicates was isolated using Trizol reagent (Invitrogen) as described ^52^.

### Cell viability analysis

Cells (1-2×10^3^) were seeded per well in 96-well plates (Thermo Fisher Scientific, Waltham, MA, USA) and incubated for 24 hours before treatment. Cell viability was assessed using Alamar Blue staining (Thermo Fisher Scientific).

Synergistic effects were evaluated using the BLISS model.

### Clonogenic survival assays

Cells were seeded at 2000 cells/well in 24-well plates and treated after 24 hours of culture. The cells were stained with 0.5% crystal violet in 25% methanol on day 7-14 after treatment and evaluated for colony formation as described ^52^.

### Tumorsphere formation assays

Cells (5 × 10^3^) were seeded per well in 6-well ultra-low attachment culture plates (Corning Life Sciences, Corning, NY, USA) and cultured as described ^52^. Tumorspheres with a diameter >200 μm were counted under an inverted microscope in triplicate wells.

### Mouse tumor model studies

Six-week-old nude female mice (The Jackson Laboratory; Bar Harbor, ME, USA) were injected subcutaneously in the flank with 2-5 × 10^6^ H358/RMC-6236-R cells in 100 μl of a 1:1 solution of medium and Matrigel (BD Biosciences) as described ^52^. Mice were pair-matched into treatment groups when the mean tumor volumes reached 100-150 mm^3^. Tumor measurements and body weights were recorded every 3-4 days. Mice were sacrificed when tumors reached >1000 mm^3^ as calculated by the formula: (width)^2^ x length/2. These studies were performed in accordance with ethical regulations required for approval by the Dana-Farber Cancer Institute Animal Care and Use Committee (IACUC) under protocol 03-029.

### Statistical analysis

Each experiment was performed at least three times. Data are expressed as the mean±SD. The unpaired Mann-Whitney U test was used to determine differences between means of groups.

Asterisks represent *P ≤ 0.05, **P ≤ 0.01, ***P ≤ 0.001, ****P ≤ 0.0001 with CI = 95%.

## Data Availability

The RNA-seq data have been deposited in the Gene Expression Omnibus under accession numbers GSE315052 and GSE324125. scRNA-seq data of KL2 tumors was obtained from GEO GSE270541 ^8^. Additional raw data is available from the corresponding author upon reasonable request.

### Acknowledgements

Research reported in this publication was supported by the National Cancer Institute of the National Institutes of Health under grant numbers CA97098, CA282437, CA289134 awarded to DK. This project has also been supported through the National Cancer Institute Experimental Therapeutics Program (NExT).

## Author Contributions

Conceptualization: ST, NH, DK; Methodology: ST, NH, KN, MM, AT, TY, TT, MDL; Investigation: ST, NH, KN, MM, AT, TY, TT, MDL; Writing – original draft: DK; Writing – review & editing: ST, NH, KN, MM, AT, TY, TT, MDL, DK; Supervision: DK; Funding acquisition: DK.

## Supplemental Figure Legends

**Supplemental Figure S1. Targeting RAS(ON) with TCIs induces M1C expression. A**. H358 cells treated with 100 nM daraxonrasib for the indicated days were analyzed for M1C transcripts. The results (mean±SD of 4 determinations) are expressed as relative levels compared to that obtained for control cells (assigned a value of 1). **B**. H2122 cells treated with the indicated concentrations of daraxonrasib for 48 hours were analyzed for M1C transcripts. The results (mean±SD of 4 determinations) are expressed as relative levels compared to that obtained for control cells (assigned a value of 1). **C**. Lysates from MGH1112 cells treated with 1 μM daraxonrasib for 48 hours were immunoblotted with antibodies against the indicated proteins. **D and E**. H358 (**D**) and H2122 (**E**) cells treated with the indicated concentrations of RMC-7977 for 48 hours were analyzed for M1C transcripts. The results (mean±SD of 4 determinations) are expressed as relative levels compared to that obtained for control cells (assigned a value of 1). **F and G**. H358 cells treated with the indicated concentrations of RMC-7977 and GO-203 (**F**) or RMC-7977 and C-11 (**G**) for 48 hours were analyzed for viability by Alamar blue staining. Indicated are the combination indices determined using Bliss scores. **H**. Lysates from H358-SR cells treated with 1 μM daraxonrasib for 48 hours were immunoblotted with antibodies against the indicated proteins. **I**. H358-SR cells treated with the indicated concentrations of daraxonrasib and C-11 for 48 hours were analyzed for viability by Alamar blue staining. Indicated are the combination indices determined using Bliss scores.

**Supplemental Figure S2. M1C confers resistance to RAS(ON) TCIs. A**. H358 and H358/DAR-R cells treated with the indicated concentrations of daraxonrasib for 48 hours were analyzed for viability by Alamar blue staining. Indicated are the IC50 values. **B**. H358/DAR-R/tet-CshRNA and H358/DAR-R/tet-MUC1shRNA cells treated with vehicle or DOX for 7 days were analyzed for M1C transcripts. The results (mean±SD of 4 determinations) are expressed as relative levels compared to that obtained for H358 cells (assigned a value of 1). **C**. H358 and H358/RMC-7977-R cells treated with the indicated concentrations of RMC-7977 for 48 hours were analyzed for viability by Alamar blue staining. Indicated are the IC50 values. **D**. H358/RMC-7977-R/tet-CshRNA and H358/RMC-7977-R/tet-MUC1shRNA cells treated with vehicle or DOX for 7 days were analyzed for M1C transcripts. The results (mean±SD of 4 determinations) are expressed as relative levels compared to that obtained for H358 cells (assigned a value of 1). **E**. H358/RMC-7977-R/tet-CshRNA and H358/RMC-7977-R/tet-MUC1shRNA cells were treated with vehicle or DOX for 7 days. The cells were then incubated with 100 nM RMC-7977 for 2 days and analyzed for colony formation. Shown are representative photomicrographs of stained colonies. The results (mean±SD of three determinations) are expressed as relative colony number compared to that for control cells (assigned a value of 1). **F and G**. H358/RMC-7977-R cells treated with the indicated concentrations of RMC-7977 and GO-203 (**F**) or RMC-7977 and M1C-11 (**G**) for 48 hours were analyzed for viability by Alamar blue staining. Indicated are the combination indices determined using Bliss scores. **H**. H2122 and H2122/DAR-R cells treated with the indicated concentrations of daraxonrasib for 48 hours were analyzed for viability by Alamar blue staining. Indicated are the IC50 values. **I**. MGH1112 and MGH1112/DAR-R cells treated with the indicated concentrations of daraxonrasib for 48 hours were analyzed for viability by Alamar blue staining. Indicated are the IC50 values. **J and K**. MGH1112/DAR-R cells treated with the indicated concentrations of daraxonrasib and GO-203 (**J**) or daraxonrasib and C-11 (**K**) for 48 hours were analyzed for viability by Alamar blue staining. Indicated is the combination index as determined using Bliss scores.

**Supplemental Figure S3. M1C localizes to cell membrane condensates in daraxonrasib-resistant cells. A**. Confocal IF imaging of H2122/DAR-R cells showing localization of M1C to condensates. Scale bar, 5 µm. **B**. Confocal IF imaging of H358/DAR-R cells left untreated and treated with 5 μM 1,6-hexanediol for 48 hours showing suppression of puncta. Scale bar, 5 µm. **C**. Confocal IF imaging of H358/DAR-R cells showing colocalization of M1C with the cell membrane Na+/K+ ATPase alpha 1 marker. Scale bar, 5 µm. **D**. H358/DAR-R cells treated with 5 μM GO-203 and 5 μM M1C-11 for 48 hours were analyzed for SHP2 transcripts. The results (mean±SD of 4 determinations) are expressed as relative levels compared to that obtained for control cells (assigned a value of 1). **E and F**. H358/DAR-R cells treated with the indicated concentrations of daraxonrasib and TNO155 (**E**) or daraxonrasib and SHP099 (**F**) for 48 hours were analyzed for viability by Alamar blue staining. Indicated are the combination indices determined using Bliss scores.

**Supplemental Figure S4. M1C activates the OSM/gp130/STAT3 axis in draxonrasib resistance. A**. RNA-seq datasets from H358/DAR-R vs H358 cells were analyzed using the HALLMARK INTERFERON ALPHA and HALLMARK INTERFERON GAMMA RESPONSE gene signatures. **B**. Chromatin from H358/RMC-6236-R cells treated with 5 μM GO-203 or 5 μM M1C-11 for 48 hours was immunoblotted with antibodies against the indicated proteins. **C**. H358 and H358/DAR-R cells were analyzed for gp130 transcripts. The results (mean±SD of 4 determinations) are expressed as relative levels compared to that obtained for H358 cells (assigned a value of 1). **D**. H2122/DAR-R cells treated with the indicated concentrations of daraxonrasib and SC144 for 48 hours were analyzed for viability by Alamar blue staining. Indicated are the combination indices determined using Bliss scores. **E**. H358/DAR-R cells expressing a CshRNA, STAT3shRNA#1 or STAT3shRNA#2 were analyzed for STAT3 transcripts. The results (mean±SD of 4 determinations) are expressed as relative levels compared to that obtained for CshRNA cells (assigned a value of 1). **F and G**. H358/DAR-R cells treated with 5 μM SC144 for 48 hours were analyzed for SHP2 transcripts (**F**). The results (mean±SD of 4 determinations) are expressed as relative levels compared to that obtained for control cells (assigned a value of 1). Lysates were immunoblotted with antibodies against the indicated proteins (**G**). **H**. Lysates from H358/DAR-R cells expressing a CshRNA, STAT3shRNA#1 or STAT3shRNA#2 were immunoblotted with antibodies against the indicated proteins.

**Supplemental Figure S5. M1C induces EMT and ALDH1A expression in daraxonrasib-resistance. A**. Lysates from H358 and H358/DAR-R cells were immunoblotted with antibodies against the indicated proteins. **B**. Lysates from H358/DAR-R cells treated with 5 μM GO-203 and 5 μM M1C-11 for 48 hours were immunoblotted with antibodies against the indicated proteins. **C**. Morphology of H358 and H358/DAR-R cells as analyzed by microscopy. **D**. Lysates from H2122/DAR-R/CshRNA and H2122/DAR-R/NF-κBshRNA cells were immunoblotted with antibodies against the indicated proteins. **E**. H2122/DAR-R/CshRNA and H2122/DAR-R/NF-κBshRNA cells treated with the indicated concentrations of daraxonrasib for 48 hours were analyzed for viability by Alamar blue staining. Indicated are the IC50 values. **F**. H2122 and H2122/DAR-R cells were analyzed for ALDH1A1 transcripts. The results (mean±SD of 4 determinations) are expressed as relative levels compared to that obtained for H2122 cells (assigned a value of 1). **G**. MGH1112 and MGH1112/DAR-R cells were analyzed for ALDH1A1 transcripts. The results (mean±SD of 4 determinations) are expressed as relative levels compared to that obtained for MGH1112 cells (assigned a value of 1).

**Supplemental Figure S6. M1C is a target for treatment of daraxonrasib-resistant cells. A and B**. H2122/DAR-R (**A**) and MGH1112/DAR-R (**B**) cells treated with 5 μM C-11 for 48 hours were analyzed for tumorsphere formation. Shown are representative photomicrographs of tumorspheres (left). The results (mean±SD of three determinations) are expressed as relative tumorsphere formation compared to that for vehicle treated cells (assigned a value of 1) (right). **C**. H358 and H358/DAR-R cells treated with the indicated concentrations of M1C ADC for 7 days were analyzed for cell viability by Alamar Blue staining. The results (mean±SD of six determinations) are expressed as relative cell number (% control) compared to that for untreated cells. Indicated are the M1C ADC IC50 values. **D**. Mean body weight changes of H358/DAR-R tumor bearing mice treated with vehicle or M1C ADC. The results are expressed as the mean ratio of relative body weight for which the SEMs were <10%.

**Supplemental Figure S7. M1C-induced resistance to daraxonrasib and sotorasib is conferred by distinct mechanisms. A**. RNA-seq datasets from H358/DAR-R vs H358-SR cells were analyzed using the HALLMARK INTERFERON ALPHA and HALLMARK INTERFERON GAMMA RESPONSE gene signatures. **B**. RNA-seq datasets from H358/DAR-R vs H358-SR cells were analyzed using the HALLMARK E2F TARGETS and HALLMARK G2M CHECKPOINT gene signatures. **C**. H358/DAR-R and H358-SR cells were analyzed for ALDH1A1 and ALDH1A3 transcripts. The results (mean±SD of 4 determinations) are expressed as relative levels compared to that obtained for H358-SR cells (assigned a value of 1). **D**. Comparison of the daraxonrasib and sotorasib resistant phenotypes.

**Supplemental Table S1.**
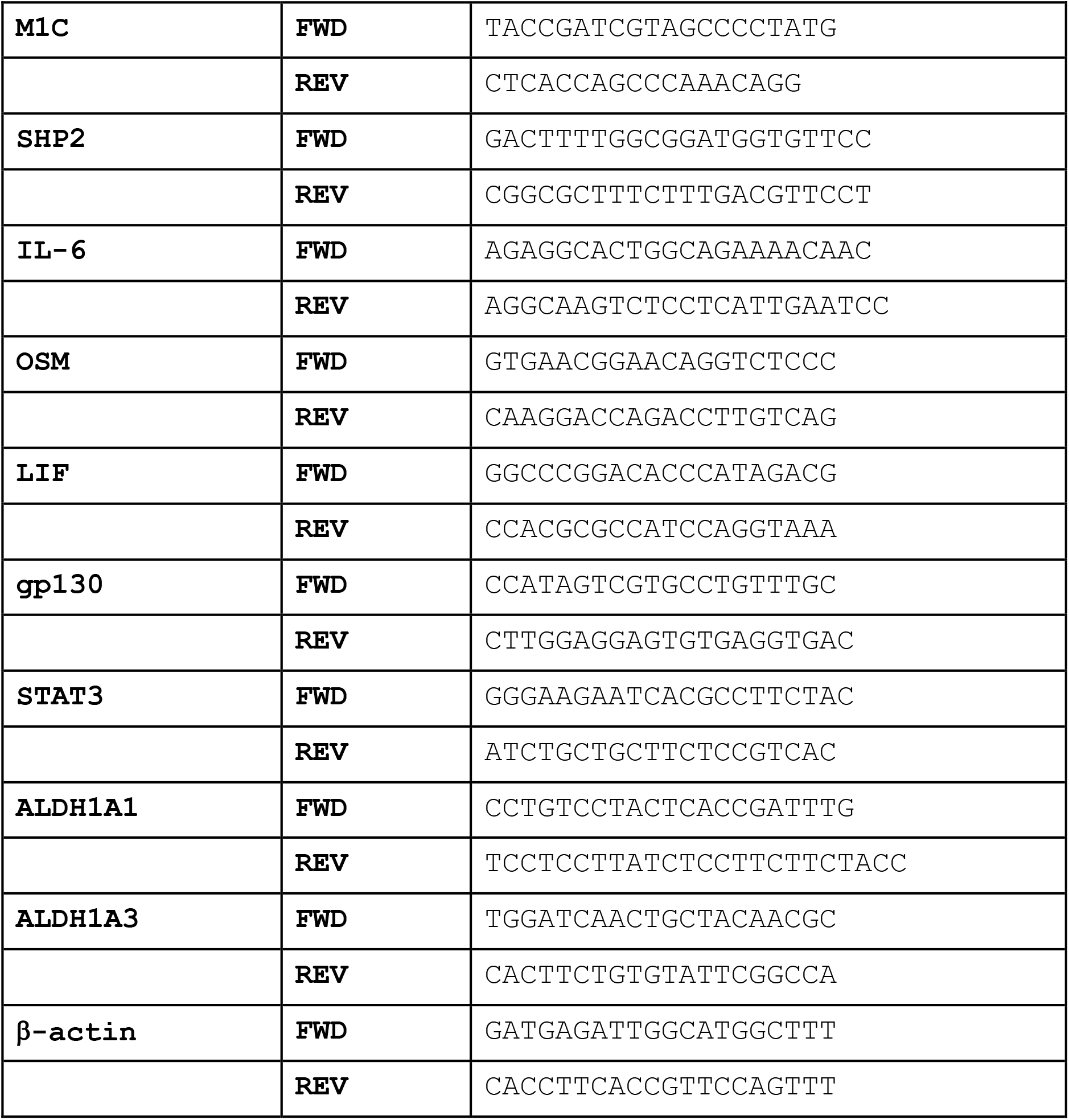
Primers used for qRT-PCR analysis.

