## Supplementary Figures for "M1C IS NECESSARY FOR DARAXONRASIB RESISTANCE OF NSCLC KRAS(G12C) MUTANT CELLS"

### Supplementary Figure 1

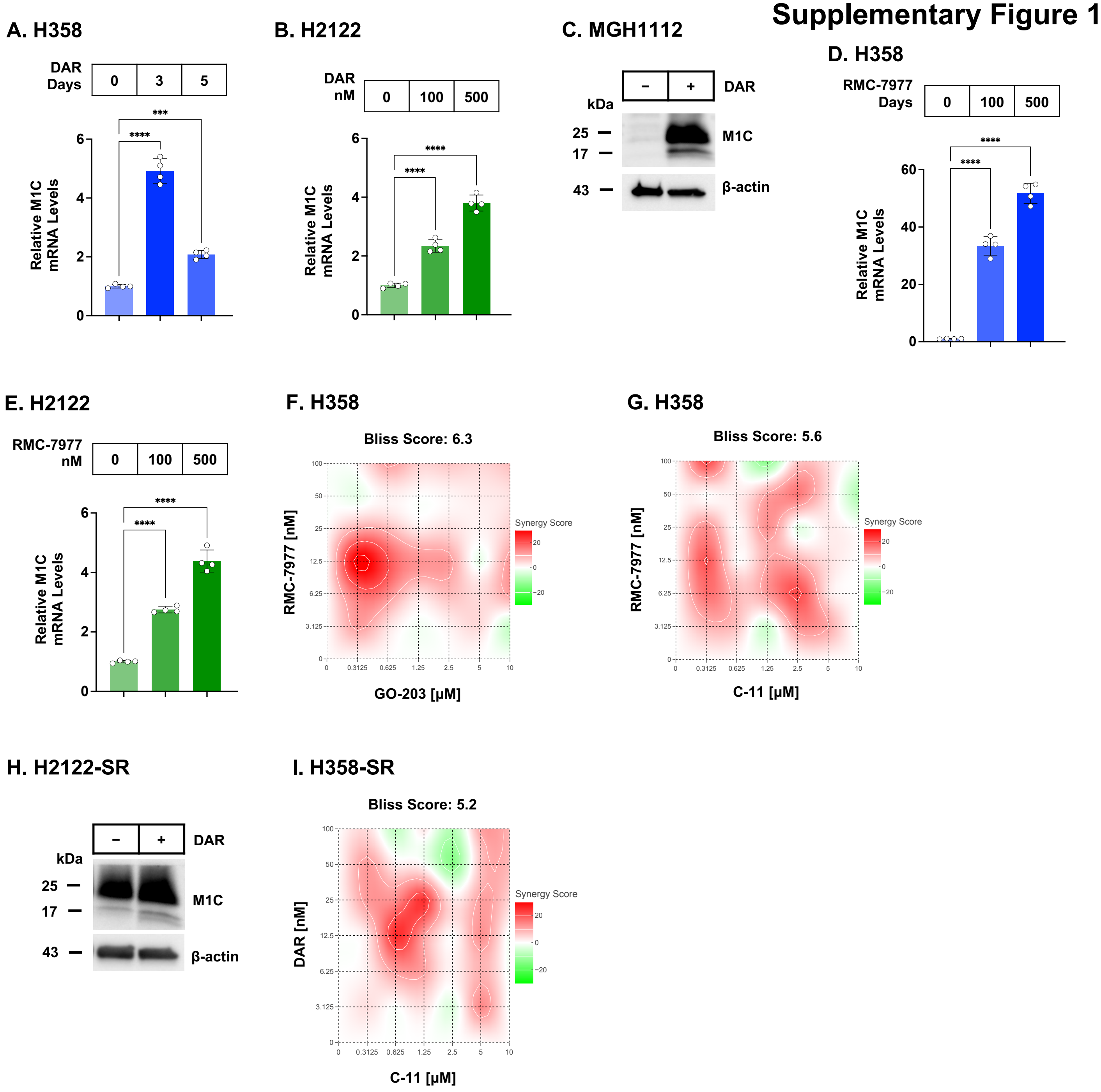

### Supplementary Figure 2

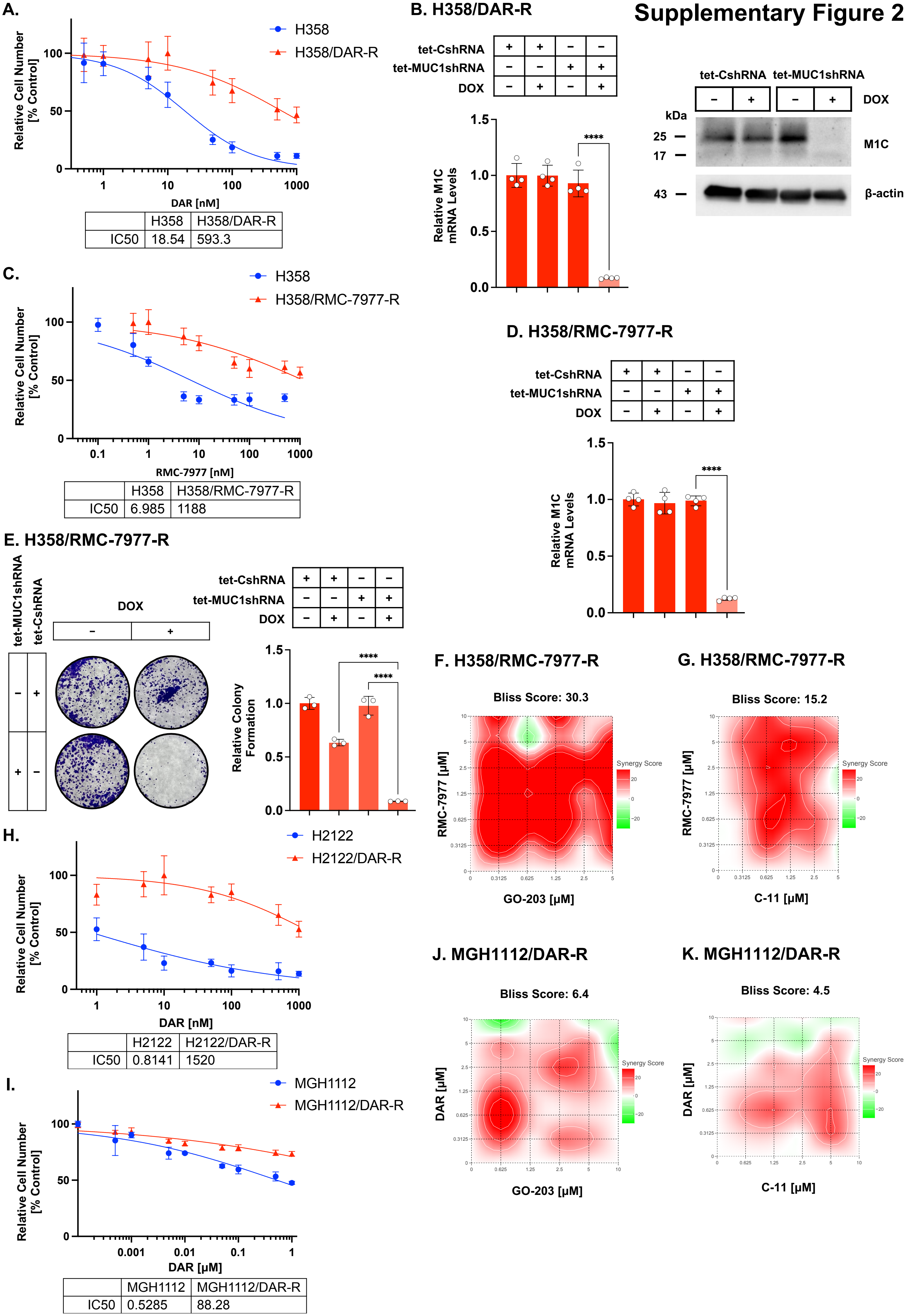


| | H358 | H358/DAR-R |
|---|---|---|
| IC50 | 18.54 | 593.3 |

| | H358 | H358/RMC-7977-R |
|---|---|---|
| IC50 | 6.985 | 1188 |

| | H2122 | H2122/DAR-R |
|---|---|---|
| IC50 | 0.8141 | 1520 |

| | MGH1112 | MGH1112/DAR-R |
|---|---|---|
| IC50 | 0.5285 | 88.28 |

### Supplementary Figure 3

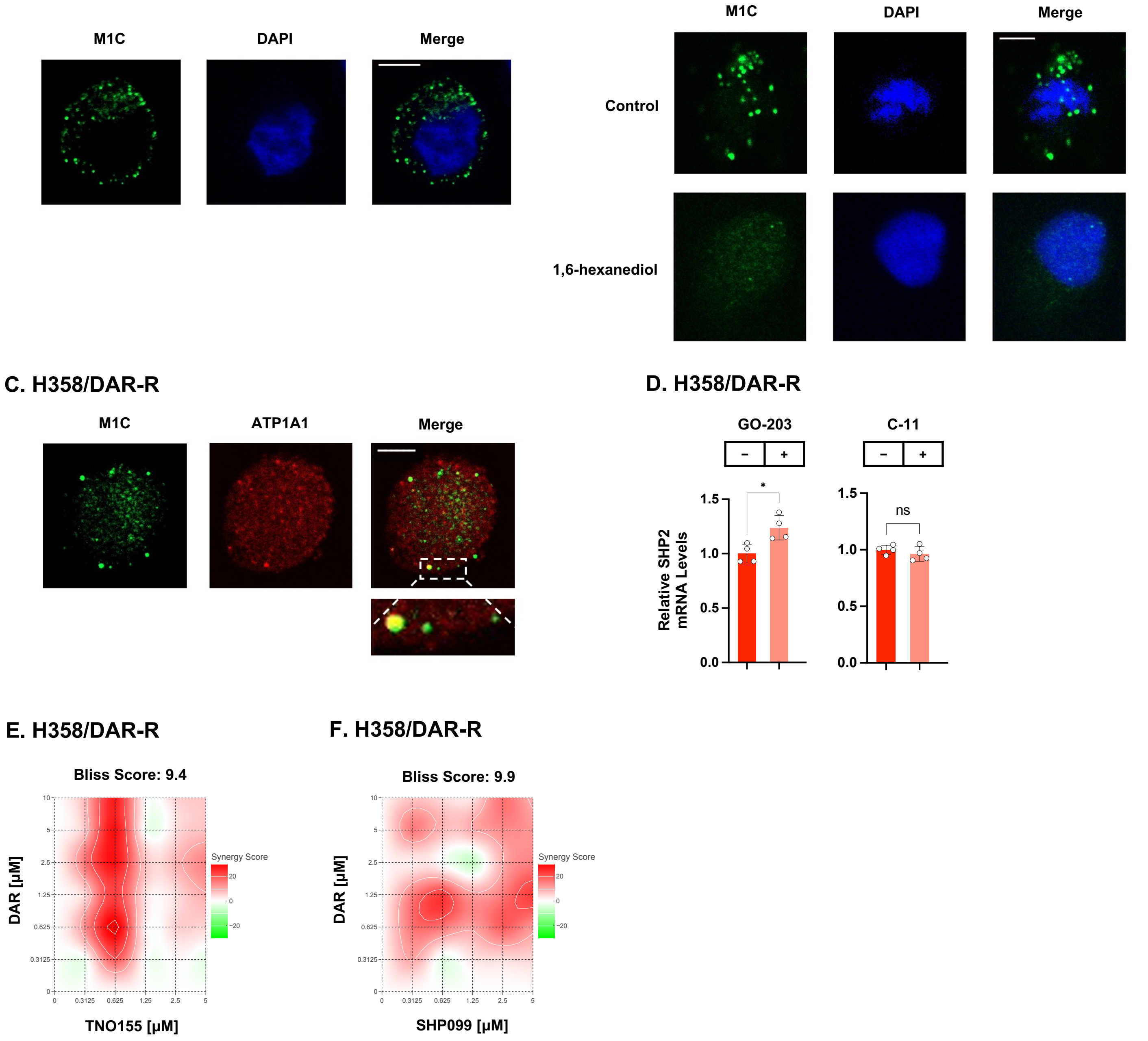

### Supplementary Figure 4

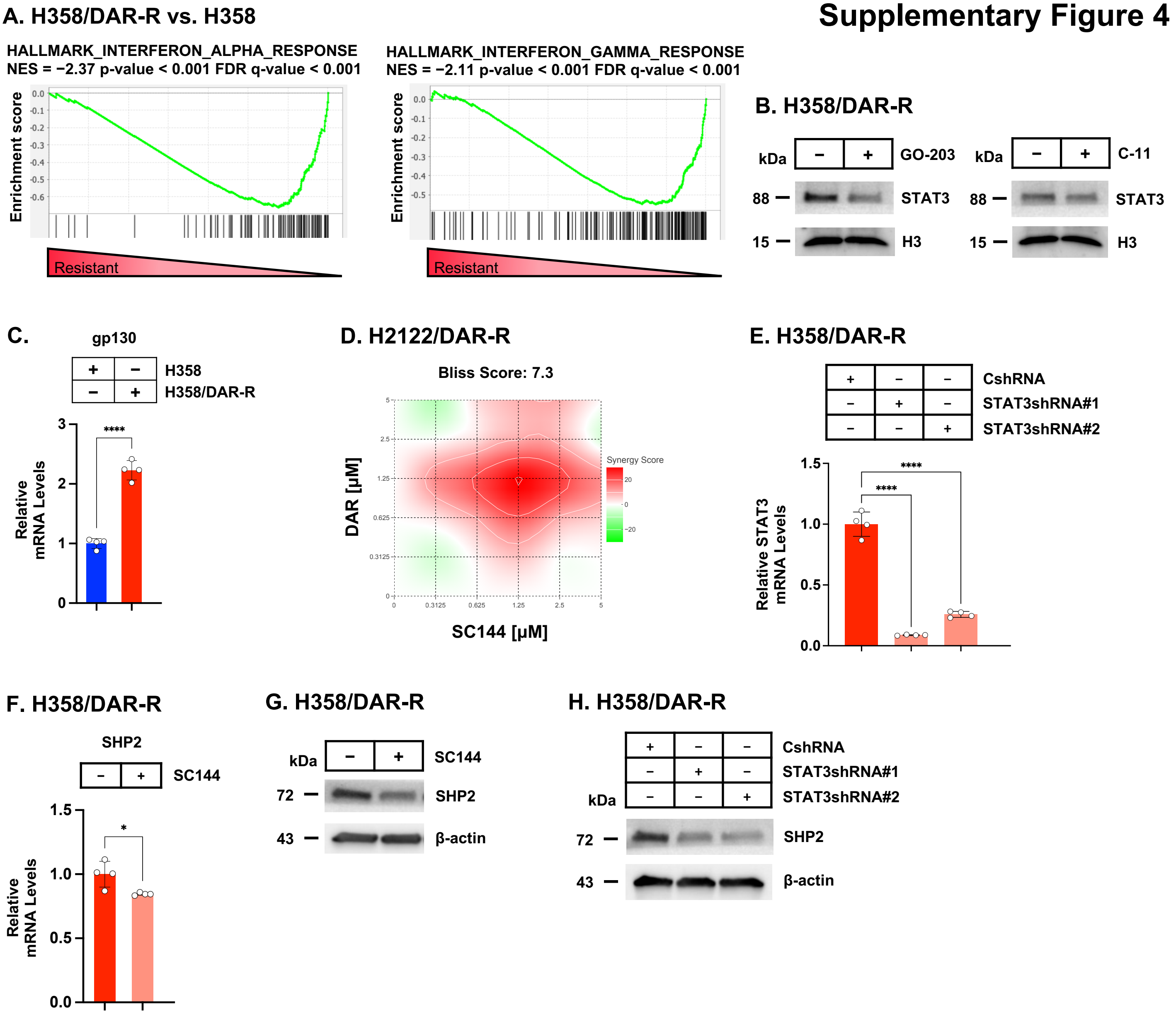

### Supplementary Figure 5

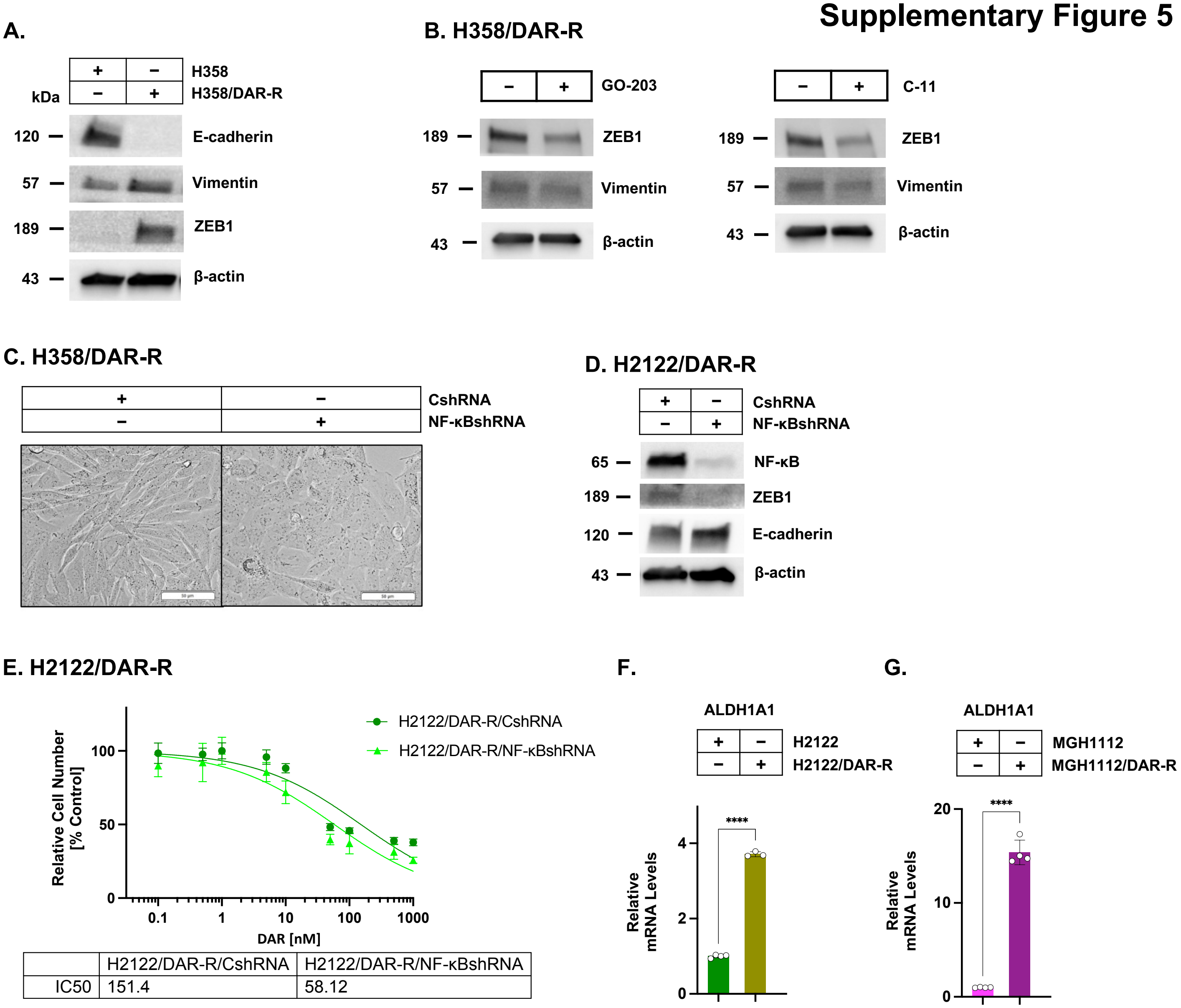


| | H2122/DAR-R/CshRNA | H2122/DAR-R/NF-κBshRNA |
|---|---|---|
| IC50 | 151.4 | 58.12 |

### Supplementary Figure 6

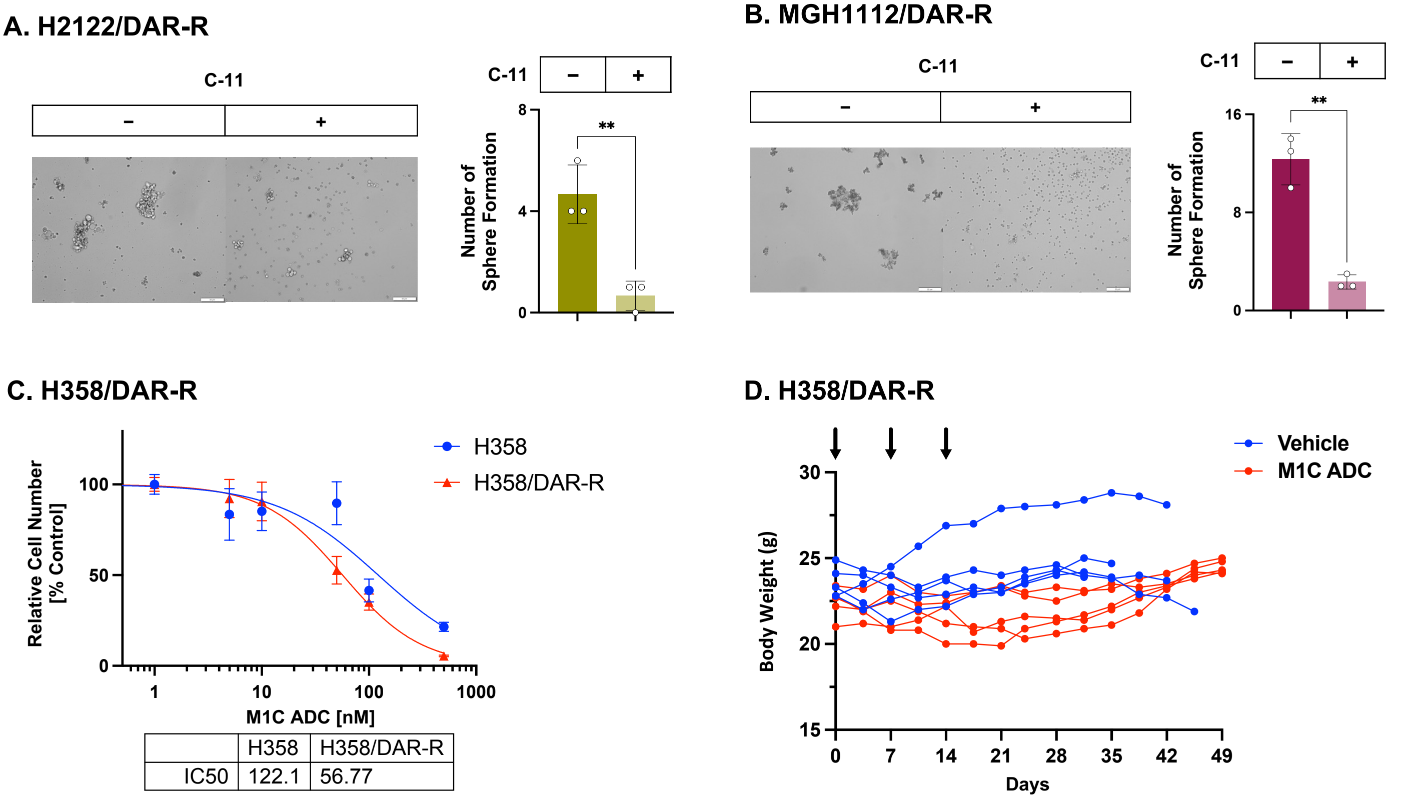


| | H358 | H358/DAR-R |
|---|---|---|
| IC50 | 122.1 | 56.77 |

### Supplementary Figure 7

#### A. H358/DAR-R vs. H358-SR

HALLMARK_INTERFERON_ALPHA_RESPONSE
NES = −2.18 p-value < 0.001 FDR q-value < 0.001

HALLMARK_INTERFERON_GAMMA_RESPONSE
NES = −2.08 p-value < 0.001 FDR q-value < 0.001

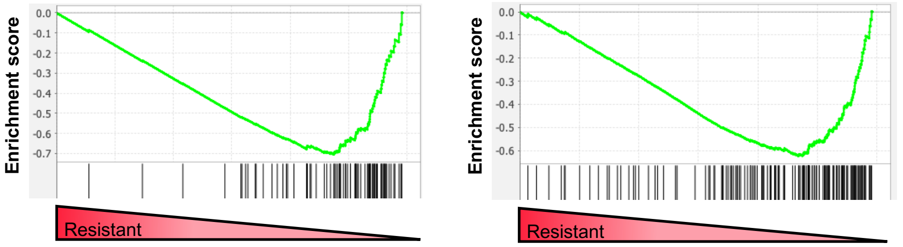


#### B. H358/DAR-R vs. H358-SR

HALLMARK_E2F_TARGETS
NES = 1.37 p-value < 0.001 FDR q-value = 0.366

HALLMARK_G2M_TARGETS
NES = 1.54 p-value < 0.001 FDR q-value = 0.203

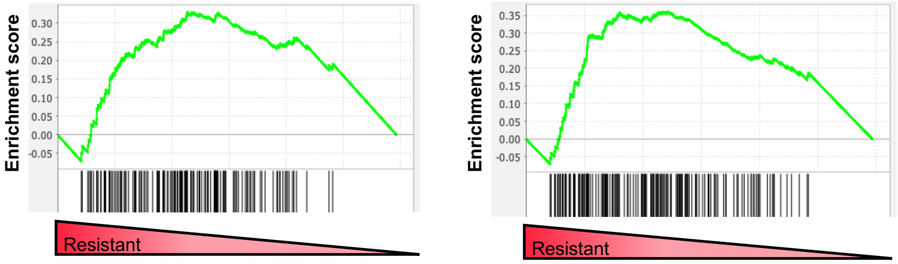


## C.

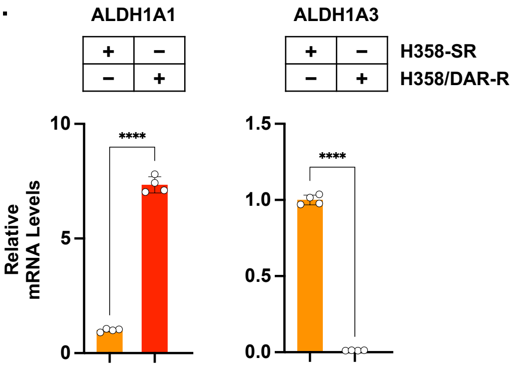


## D.

| RESISTANCE | SOTORASIB | DAR |
|---|---|---|
| **Transcriptome** | | |
| IFN | + | − |
| G2M | − | + |

| | | |
|---|---|---|
| **Upregulation** | | |
| OSM | − | + |
| gp130 | − | + |
| STAT1 | + | − |
| STAT3 | − | + |
| EMT | + | + |
